# Two distinct bacterial biofilm components trigger metamorphosis in the colonial hydrozoan *Hydractinia echinata*

**DOI:** 10.1101/2019.12.23.887182

**Authors:** Huijuan Guo, Maja Rischer, Martin Westermann, Christine Beemelmanns

**Affiliations:** Leibniz Institute for Natural Product Research and Infection Biology – Hans Knöll Institute, Beutenbergstraße 11a, D-07745 Jena, Germany; Electron Microscopy Centre, Friedrich Schiller University Jena, Ziegelmühlenweg 1, D-07743 Jena, Germany

**Author notes:** contributed equally (Huijuan Guo, Maja Rischer). **Author Contributions:** Conceptualization: M.R, H.G, M.W., C.B.; Methodology: M.R, H.G, M.W., C.B.; Investigations: M.R, H.G, M.W., C.B.; Resources: M.W., C.B.; Writing – Original Draft: M.R, H.G, C.B. Supervision and Funding Acquisition: M.W., C.B.

**Keywords:** *Hydractinia*, *Pseudoalteromonas*, metamorphosis, phospholipids, polysaccharides

## Abstract

In the marine environment bacterial-induced metamorphosis of larvae is a widespread cross-kingdom communication phenomenon and critical for the persistence of many marine invertebrates. However, the identities of most inducing bacterial signals and the underlying cellular mechanisms remain enigmatic. Larvae of *Hydractinia echinata* provide an excellent model for investigating bacteria-stimulated settlement as they transform upon detection of the signal into the colonial adult stage within 24 h. Although *H. echinata* served as cell biological model system for decades, the influence of bacterial signals on the morphogenic transition remained largely unexplored. Using a bioassay-guided analysis, we first identified that specific bacterial (lyso)phospholipids, naturally present in bacterial biofilms, elicit metamorphosis in *Hydractinia* larvae in a dose-response matter. In particular, lysophospholipids as single compounds or in combinations at 50 µM concentrations induced metamorphosis in up to 50% of all larvae phospholipid within 48 h. By using fluorescence-labeled bacterial phospholipids, we demonstrated their incorporation into the larval membranes, where interactions with internal signaling cascades could occur. In addition, two structurally distinct exopolysaccharides, the newly identified Rha-Man polysaccharide from *Pseudoalteromonas* sp. P1-9 and curdlan from *Alcaligenes faecalis* caused up to 75% of all larvae to transform within 24 h. We also found that combinations of (lyso)phospholipids and curdlan induced the transformation in almost all larvae within 24 h, thereby exceeding the morphogenic activity observed for single compounds and axenic bacterial biofilms. Our results demonstrate that multiple and structurally distinct bacterial-derived metabolites converge to induce high transformation rates of *Hydractinia* larvae, which might ensure optimal habitat selection despite the general widespread occurrence of both compound classes.

**Significance Statement:** Bacterial biofilms profoundly influence the recruitment and settlement of marine invertebrates, critical steps for diverse marine processes such as coral reef formation, marine fisheries and the fouling of submerged surfaces. Yet, the complex composition of biofilms often makes it challenging to characterize the individual signals and regulatory mechanisms. Developing tractable model systems to characterize these co-evolved interactions is the key to understand fundamental processes in evolutionary biology. Here, we characterized for the first time two types of bacterial signaling molecules that induce the morphogenic transition and analyzed their abundance and combinatorial activity. This study highlights the crucial role of the converging activity of multiple bacterial signals in development-related cross-kingdom signaling.

**Areas:** Major: Chemical Biology, Microbiology, Developmental Biology

## Introduction

The radical transformation (metamorphosis) of motile larvae into the adult stage is a critical step in the life cycle of many marine species as it confers the propagation and persistence of the population in the marine ecosystem.^1^ For more than 80 years it has been recognized that chemical signals present within marine biofilms induce the settlement and metamorphosis in larvae,^2,3,4^ but the identification of these signaling molecules remains still a challenging task and until today and only very few key bacterial signals have been structurally characterized and functionally analyzed.^5,6,7^ One of the few identified bacterial signaling molecules are bromopyrroles produced by *Pseudoalteromonas* that induce metamorphosis in coral larvae,^8,9^ but larvae failed to attach to surfaces when stimulated by bromopyrroles alone and it was deduced that other, yet unidentified, chemical cues are important for the morphogenic process. Recent biochemical investigations of the bacteria-induced metamorphosis of the marine polychaete *Hydroides elegans* resulted in the identification of a phage tail-like contractile injection systems (tailocins) in *Pseudoalteromonas* species that induce settlement and metamorphosis by releasing an effector protein Mif1, which stimulates the P38 and MAPK signaling pathways.^10,11,12^ However, bacteria not capable of producing theses proteinaceous injection systems were also found to induce the transformation presumably by other, not yet identified morphogens.^13,14^ In the 1970s, Leitz and Wagner reported that a lipid-like molecule of *Pseudoalteromonas espejiana* (original name: *Alteromonas espejiana*) induces larvae transformation in *Hydractinia echinata*, an early branching metazoan lineage dating back more than 500 million years.^15^ But despite intensive studies, the bacterial signals causing *Hydractinia* larvae to metamorphose have remained elusive. Instead, metamorphosis of *Hydractinia* was artificially induced using high salt concentrations (CsCl).^16^ While highly effective, artificial induction caused phenotypical and developmental differences in *Hydractinia* development compared to bacterial induction.^17^

To gain insights into the bacteria-*Hydractinia* dependencies we set out to characterize the bacterial morphogens from associated *Pseudoalteromonas* species and phylogenetically related bacterial lineages using a bioassay-guided isolation approach. Here, we report on the identification and structure-activity studies of two types of bacterial metabolites that alone and in combination stimulated *H. echinata* larvae to metamorphose into the adult stage. Our results also suggest that OMVs and extracellular matrix play an essential role in the prokaryote-eukaryote signaling mechanisms, which will likely hold true for many other benthic marine invertebrate animals.

## Results

### Bioassay-guided identification of morphogenic bacteria

Fertile *H. echinata* polyps release their eggs or sperm in a light-controlled process, whereupon the eggs are fertilized. Within 72 h, embryos develop into a planula larva after passing through several stages, which are able to move through cilia. Active searching behavior allows the larva to explore surfaces and upon detection of bacterial signals develop into a primary polyp within 24 h. To describe the herein applied larvae-based metamorphosis assay more precisely, four different states were defined (Figure 1):^15,16^ (1) no induction: larvae continue swimming or were reversible attachment to surface; (2) settlement (SET): irreversible attachment to surface (oral ending), contraction along their oral-aboral axis, flat disc with a small tip in the centre, but not yet fully elongated to primary polyp; (3) metamorphosis (MET): full transformation into functional primary polyp and visible formation of tentacles, a hypostome and stolons; (4) dead larvae with lysed body parts.

**Figure 1.**
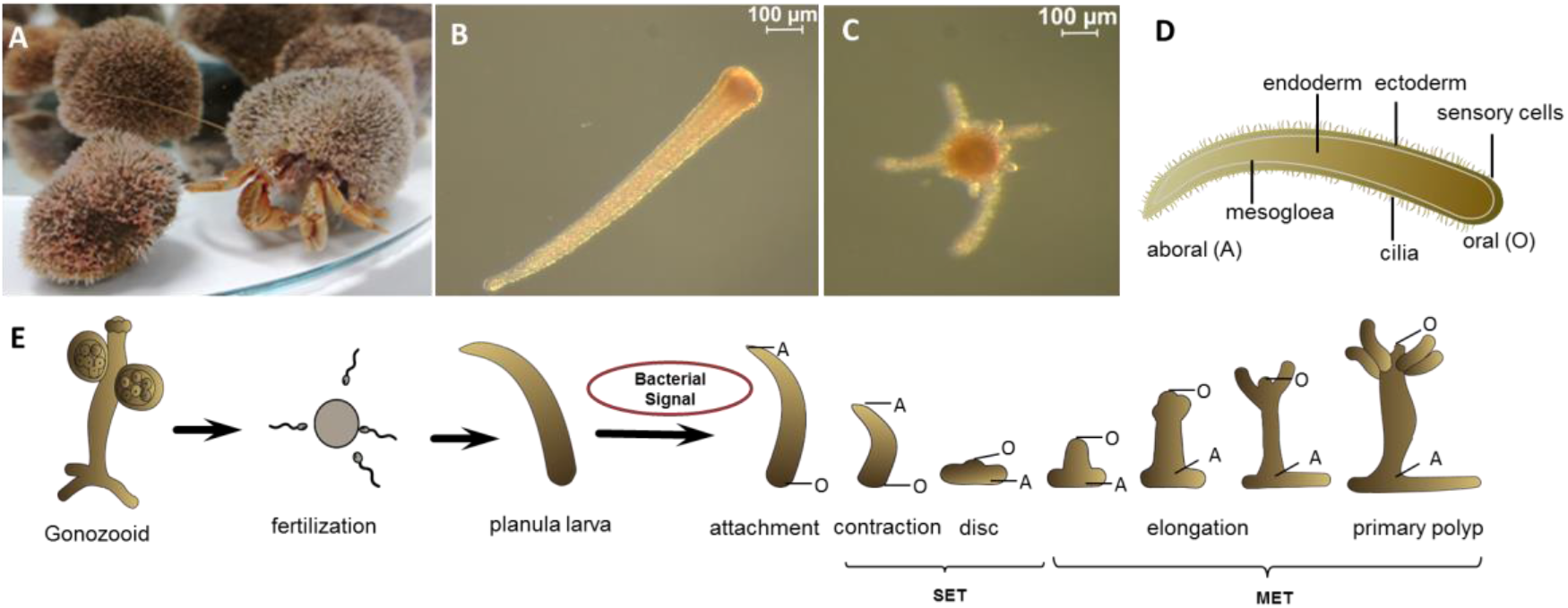
A) *Hydractinia echinata* colonizing the shell inhabited by a hermit crab (*Pagurus* sp.); B)-C) Microscopic image of *H. echinata* B) planula larva and C) primary polyp. D) Sketch of a larvae. E) Generalized sketch of the life cycle of *Hydractinia*.

Although early reports indicated that bacterial isolates belonging to different bacterial lineages (e.g. *Pseudoalteromonas* spp., *Oceanospirillum spp*., *Staphylococcus aureus, Pseudomonas* spp, *Serratia marinorubra*) are able to induce metamorphosis,^18^ many of the earlier reported strains were neither available nor phylogenetically correctly assigned. Thus, we evaluated the metamorphosis inducing capacity of twenty representative and phylogenetically characterized bacterial strains isolated from *Hydractinia*-covered shells,^19,20,21^ one phylogenetically-related coral associated strain *Pseudoalteromonas sp*. PS5,^22^ and eight bacterial type strains using a previously reported mono-species biofilm-based metamorphosis assay.^23^ As negative control, larvae were kept in artificial sea water, while larvae treatment with CsCl (final concentration of 6 mM) was used as positive control. *Hydractinia* larvae responded to the presence of CsCl in a highly synchronized and morphological distinct manner tolerating the continuous presence of 6 mM CsCl without any signs of abnormal transformation. In contrast, we observed non-synchronized induction events in our colony-based assays, which presumably results from the complex nature of bacterial biofilms and the spatial fluctuation of the yet unidentified bacterial signaling molecules. Thus, we determined settlement (SET) and metamorphosis (MET) rates after 24 and 48 h. As shown in Figure 2, three bacterial strains (*Pseudoalteromonas* sp. P1-9, *Pseudoalteromonas* P1-29, *Exiguobacterium* sp. P6-7) were found to induce settlement and metamorphosis to the primary polyp within 24 h.

**Figure 2.**
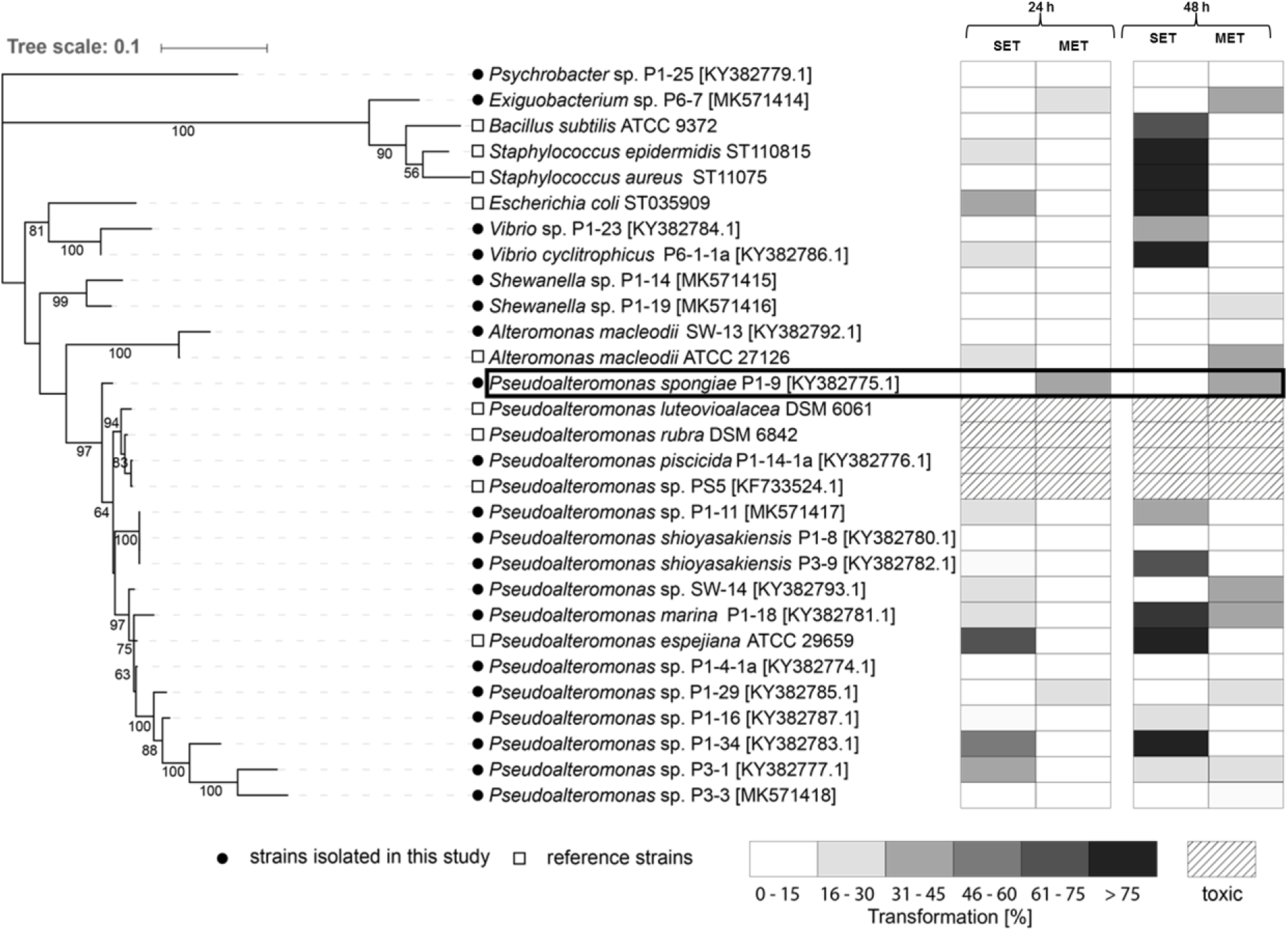
A) Left: phylogenetic tree based on 16S rRNA gene sequences of 29 tested bacterial strains. Best DNA model was generated and the robustness of interfered tree topologies was evaluated after 1000 bootstraps (> 50% are shown). Right: Heatmap depicting [%] of settled larvae (SET) and larvae that underwent metamorphosis (MET) after 24 and 48 h. Mean values of replicates are visualized as color-coded 15% step-gradient (n = 3, negative control: ASW; positive control: CsCl).

Additional five strains (*Pseudoalteromonas* sp. P1-19, *Pseudoalteromonas* sp. SW-14, *Pseudoalteromonas marina* P1-18, *Pseudoalteromonas* sp. P3-1, *Alteromonas macleodii* ATCC27126) were found to induce the settlement within the first 24 h, but subsequent formation of the primary polyp was observed only within 48 h. Up to eleven strains, including several Gram-positive reference strains, induced settlement, but neither of those strains induced the full transformation to the primary polyp causing eventually the death of the transforming larvae. Six strains were found to be non-inducing and four strains, mostly belonging to the clade of pigmented *Pseudoaltermonas* strains,^24^ caused the death of up to 100% of all larvae within 24 h. Across all tested strains, *Pseudoalteromonas* sp. P1-9 (referred from now on as P1-9) was found to induce the most robust morphogenic response and thus was selected for further detailed chemical analysis.

### Bioassay-guided identification of bacterial signals from *Pseudoalteromonas* sp. P1-9

To test if the morphogenic cue is a secreted and/or a diffusible small metabolite, methanolic solid-phase extracts (C18 cartridges) of supernatants from liquid and plate cultures of P1-9 were prepared according to previously described procedures;^23^ however, none of the metabolite extracts showed morphogenic activities. We then tested if the signal is a secreted high-molecular weight (HMW) biomolecule (e.g. protein, cell surface-associated exopolysaccharide (EPS), (EPS)) and/or part of the bacterial membranes (Figure 3, Figure S2-S6). Size exclusion separation of concentrated cell membrane components resulted in the isolation of a HMW fraction (> 30 kDa) that showed morphogenic activity in a dose-response manner causing up to 80% of all larvae to metamorphose (entry 5). In contrast, low-molecular weight fractions (< 5 kDa) revealed only moderate activities causing less than 40% of all larvae to transform (entry 6). To test if the signal is an integral part of outer membrane vesicles (OMVs),^25,26,27^ sterile-filtered culture supernatant was separated by ultracentrifugation and, indeed, the resulting OMV-enriched precipitate was found to induce metamorphosis in more than 80% of all larvae within 24 h (entry 7). Similar results were obtained with enriched exopolysaccharide fractions (entry 8, Figure S10). To confirm that the active fractions contain OMVs and/or biopolymers fractions and cultures were subjected to electron microscopy (EM) imaging. While scanning electron microscopy pictures indicated that *Pseudoalteromonas* sp. P1-9 actively releases OMVs (Figure 4A), (cryo)TEM imaging of bacterial cells obtained from liquid and plate growth showed entire cells, minicells and cells proliferating what appear to be outer membrane vesicles (100-300 nm) as well as high abundances of extracellular EPS-like fibers (Figure 4B,C). At this stage, we reasoned that the morphogenic cues might be part of the outer cell membrane and presumably of high molecular weight.

**Figure 3.**
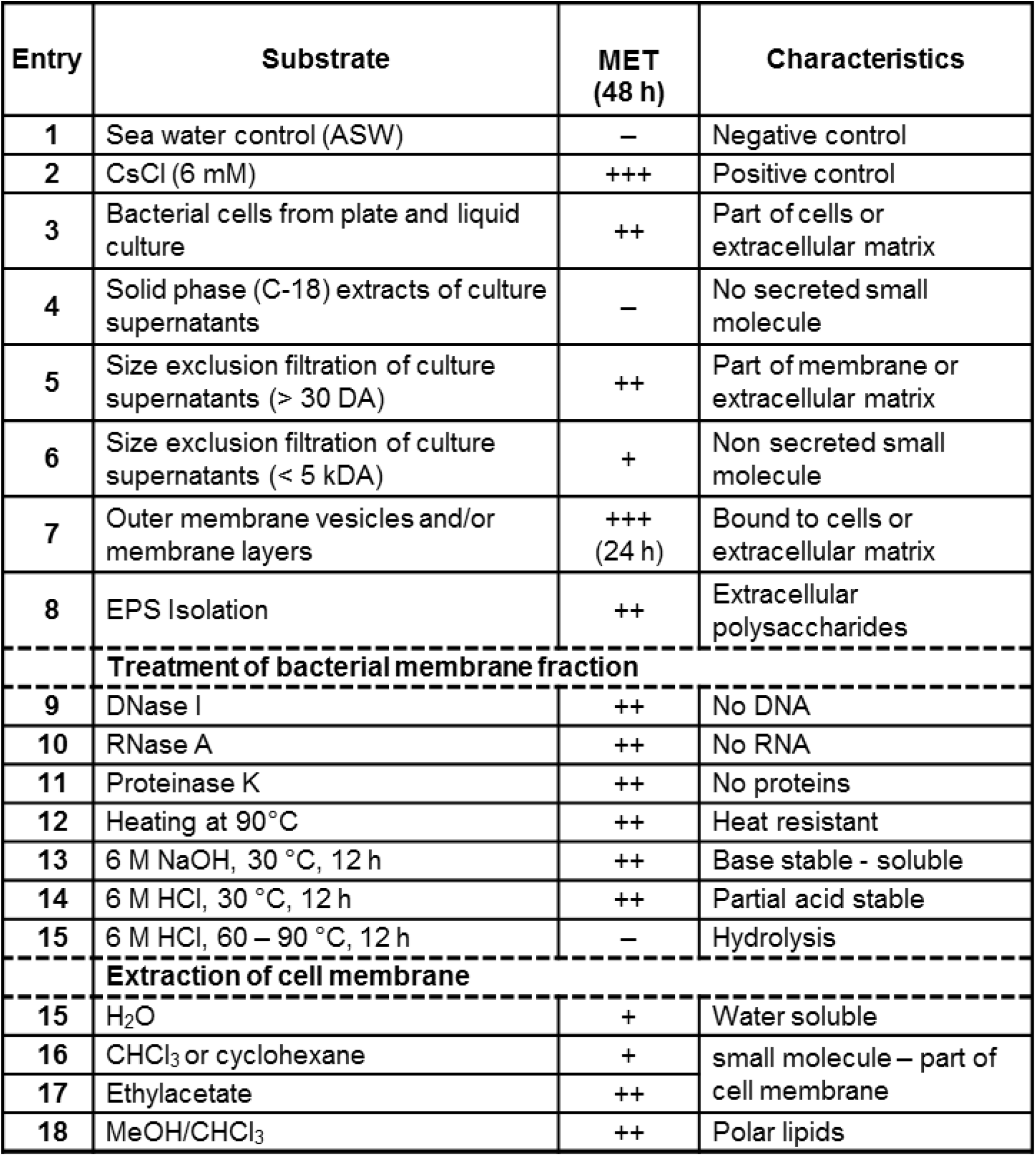
Morphogenesis assay of biosamples (n = 3) derived from *Pseudoalteromonas spongiae* P1-9. percentage of metamorphosis was evaluated after 48 h (- (no induction); + (< 40% of all larvae metamorphosed); ++ (between 40% and 80% if all larvae metamorphosed), +++ (more than 80% of all larvae metamorphosed).

**Figure 4.**
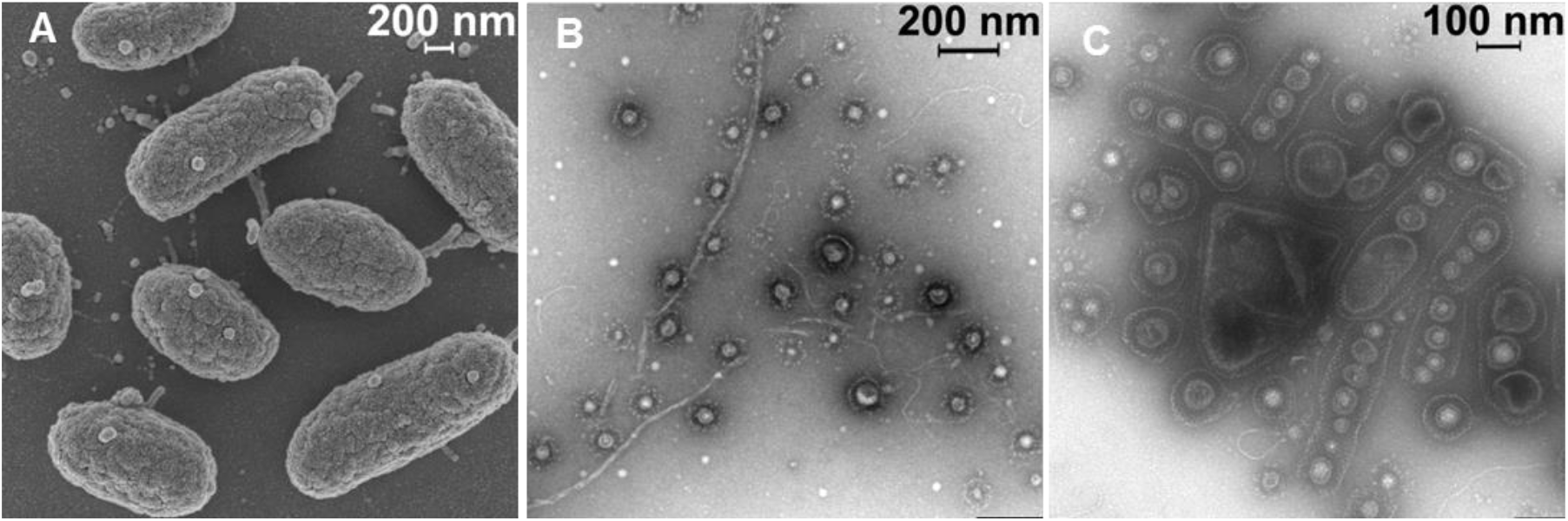
A) Scanning electron microscopy of single cells of P1-9 obtained from a three day old liquid culture. B and C)) Negative contrast electron microscopic image of vesicles coated with S-layer like matrix and string-shaped biopolymers isolated from a three day old liquid culture of P1-9 (C) and agar plate derived P1-9 biofilm (D).

To determine its stability and chemical nature, we subjected the active HMW fractions (> 30 kDa) to enzymatic and physical treatments prior to activity tests (Figure S5-S9). As depicted in Figure 2, activity was completely retained when samples were treated with digestive enzymes such as DNase, RNase, or proteinase K, or even when heated to 96 °C for 10 min (entries 9-12). Treatment with aqueous 6 M NaOH or 6 M HCl (12 h, 30 °C) partially solubilized the active morphogen (entry 13, 14), and activity of both, soluble fraction and residue, was mostly retained after neutralization. However, treatment with 6 M HCl at higher temperatures (> 60 °C) gradually abolished the morphogenic activity of the sample. Based on these tests, we deduced that the morphogen(s) was neither a sensitive protein, nucleic acid nor an instable secreted metabolite. This time, methanolic extracts as well as aqueous extracts of enriched membrane fractions showed morphogenic activity (entry 15, 17, 18). Taken together, our results indicated that *Pseudoaltermonas* sp. P1-9 produces two different bacterial morphogens, which are chemically stable and likely associated or part of the bacterial outer membrane.

### Analysis of morphogenic phospholipids

In a next step, we applied a bioassay-guided reverse-phase column chromatography (HPLC) purification protocol to purify the methanolic extracts of enriched membrane. The resulting active HPLC fractions were analyzed by high-resolution tandem mass spectrometry (HR-MS^2^) and Global Natural Product Social Molecular Networking (GNPS)^28^ analysis, which revealed the dominant presence of phosphatidylglycerols (PGs) and phosphatidylethanolamines (PEs) and the respective lyso derivatives (LPGs, LPEs) (Figure S11-S13). Due to inherent difficulties associated with the purification of structurally closely related phospholipids, commercial derivatives with matching LC-HRMS^2^ pattern (Figure S40-S50) and ^1^H and ^13^C nuclear magnetic resonance (NMR) spectra (Figure S26-S39) as well as two fluorescence-labeled derivatives were tested in a standardized assay (Figure 5, Figure S14-S15, Table S3). Two lysophospholipids (16:0 LPG, 16:0 LPA), phosphatidylethanolamine 18:0 PE, and 1,2-dipalmitoyl-sn-glycero-3-phosphate (16:0 PA) repeatedly induced settlement and metamorphosis in 20-40% of all larvae within 48 h, while phospholipid concentrations exceeding 50 µM occasionally caused lysis of larvae (Figure 5).

**Figure 5.**
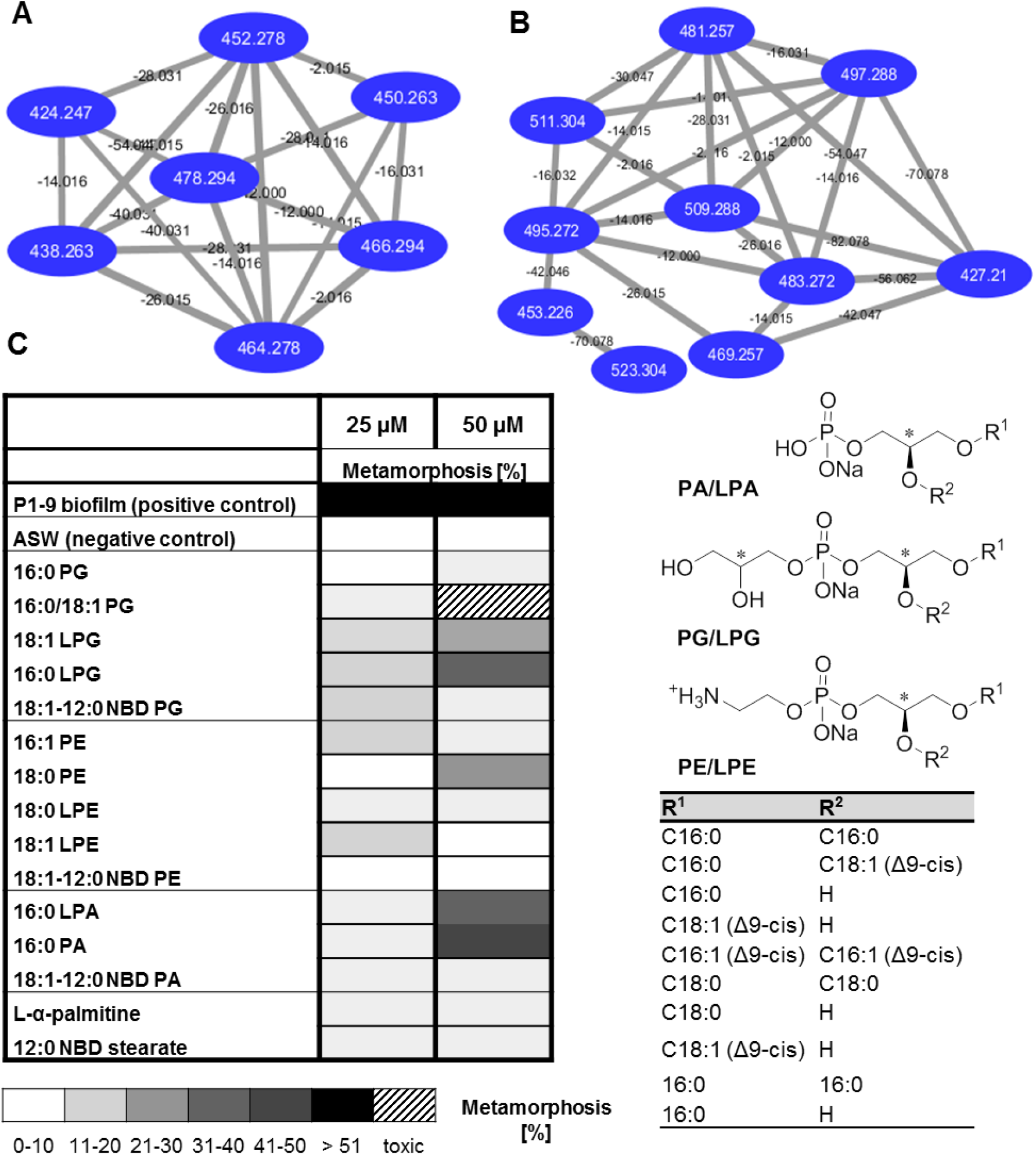
A-B) LC-HRMS^2^ based GNPS analyses of purified lipid fraction showing the MS^2^- cluster of A) LPE and B) LPG. C) Metamorphosis inducing activity of commercial phospholipids. Test were performed at 25 and 50 µM concentrations and transformation rates were determined after 24 and 48 h. Mean values of replicates are visualized as color-coded 10% step-gradient (n = 3, not shown: CsCl as positive control).

We questioned if the low solubility of (lyso)phospholipids might prevent the uptake and perception by *Hydractinia* larvae and treated competent larvae (20-30 larvae) with nitrobenzoxadiazole (NBD)-labeled (phospho)lipids (18:1-12:0 NBD PG, 18:1-12:0 NBD PE or 12-NBD stearate (Figure 6, Table S3)). Fluorescence was monitored for 24 h resulting in more than 80% fluorescence-labeled larvae, of which approx. 20% underwent transformation to the primary polyp. During metamorphosis the fluorescence signal continuously decreased, presumably due to internalization and decomposition of phospholipids during the transformation processes (Table S3). Overall, it was deduced that despite their low solubility phospholipids appear to be incorporated into the cell membranes of larvae and could interact with intercellular signaling pathways.

**Figure 6.**
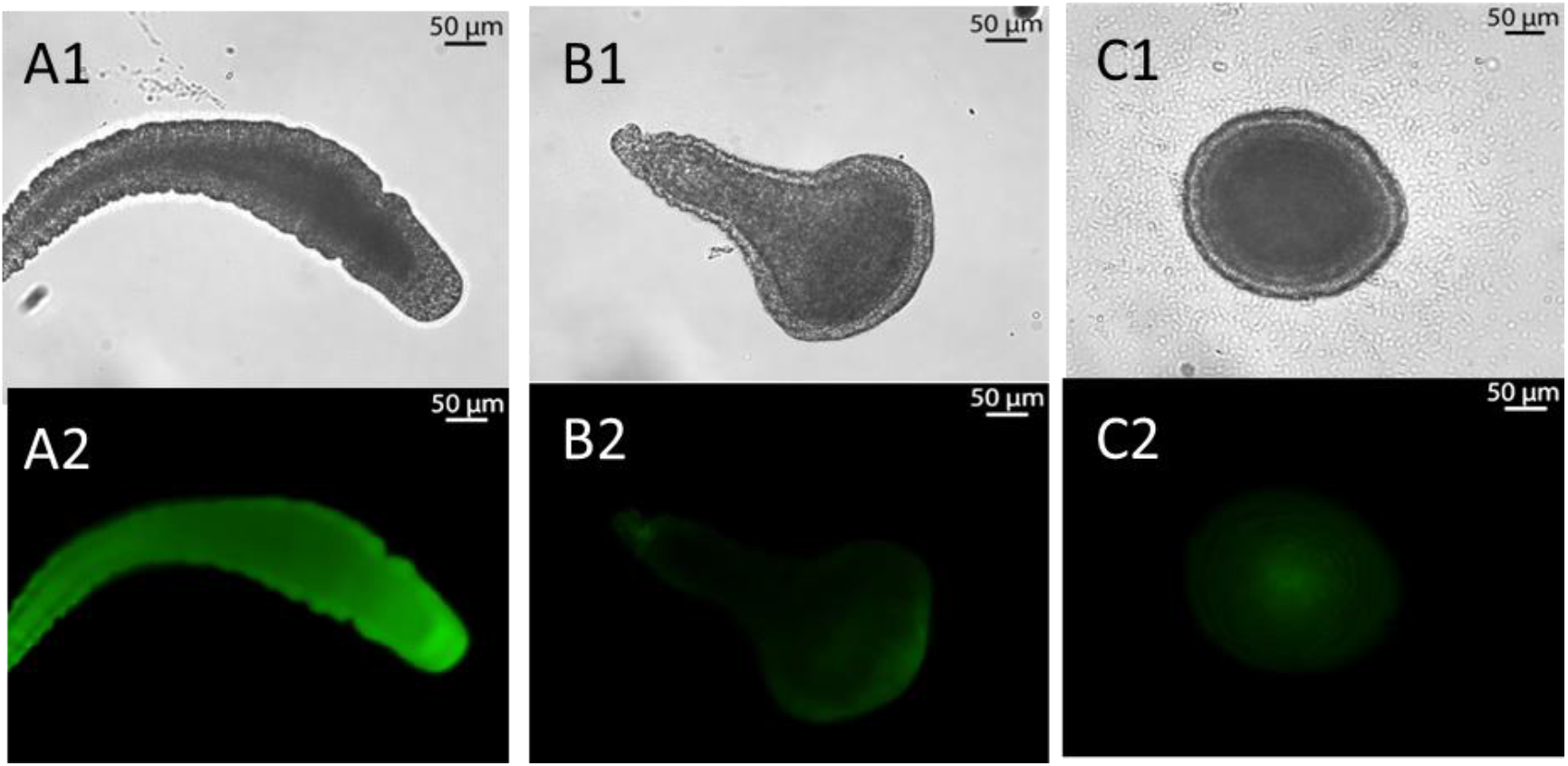
Metamorphosis assay with fluorescence-labeled phospholipids. A-C) Representative fluorescence labeling images of larvae treated with (A) 18:1-12:0-NBD PG after 5h; (B) 18:1-12:0-NBD PE after 24 h; (C) 18:1-12:0 NBD PA after 24 h (upper row: bright field, bottom row: fluorescence imaging with emission wavelength at 527 nm).

We then questioned if phospholipid combinations might exhibit synergistic, inhibitory or toxic effects and treated larvae with 1:1 combinations of two lipids (each 25 µM). As shown in Figure 7, combinations of 18:1 LPE/16:0 LPG, 18:1 LPE/18:1 LPG, 16:0 PA/16:0 PG, 16:0 LPA/16:0 LPG and 16:0 LPA/18:1 LPE, showed indeed additive if not synergistic tendencies by inducing metamorphosis in up to 50% of all larvae within 48 h (Figure 5), while others combinations appeared to have toxic effects.^29^ At this point we concluded that specific (lyso)phospholipids and combinations thereof, known to be present in bacterial cell membranes and OMVs induce metamorphosis of *Hydractinia* larvae, but are only partially reflect the observations made in assays of bacterial biofilms.

**Figure 7.**
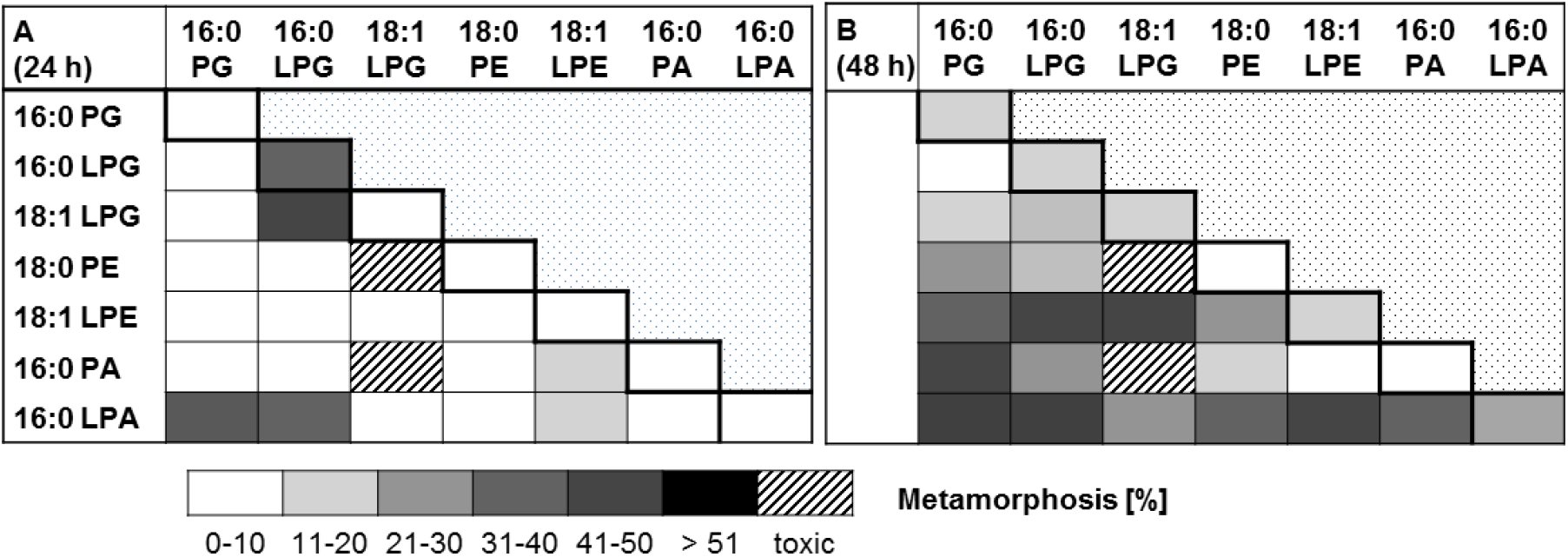
Metamorphosis assay of phospholipid combinations (25 µM of each lipid in a 1:1 combination). Transformation rates were monitored after A) 24 h and B) 48 h. Mean values of replicates are visualized as color-coded 10% step-gradient (N = 3, not shown: negative control: ASW; positive control: CsCl)

As (PL) and to a lesser extend lysophospholipids (lyso-PLs, LPLs) are species–specific phospholipids components of bacterial cell walls and outer membrane vesicles (OMVs),^30,31^ we hypothesized that the lipid composition of a bacterial strain and or biofilm will dictate the morphogenic response of larvae. To validate this hypothesis, phospholipids extracts of 15 bacterial species were analyzed by comparative HRMS^2^ and tested for their morphogenic properties (Figure 8). Indeed, lipid extracts obtained from six bacterial strains (*Vibrio* sp. P1-23, *A. macleodii*. SW13, *P. rubra, P. shioyasakiensis* P3-9, *Pseudoalteromonas* sp. P1-4-1a, *Pseudoalteromonas* sp. P1-16) induced metamorphosis to the primary polyp in 30-80% within 48 h, while lipid extracts of *Vibrio cyclitrophicus* P6-1-1a, *Pseudoalteromonas piscidida* P1-14-1a, and *P. espejiana* ATCC 29659 revealed no inducing activity. Extracts of four other strains caused cell lysis and/or larvae death, presumably due to the presence of cytotoxic secondary metabolites (violacein and/or poly-brominated pyrrole derivatives) and/or the presence of toxic phospholipid combinations.^32^

**Figure 8.**
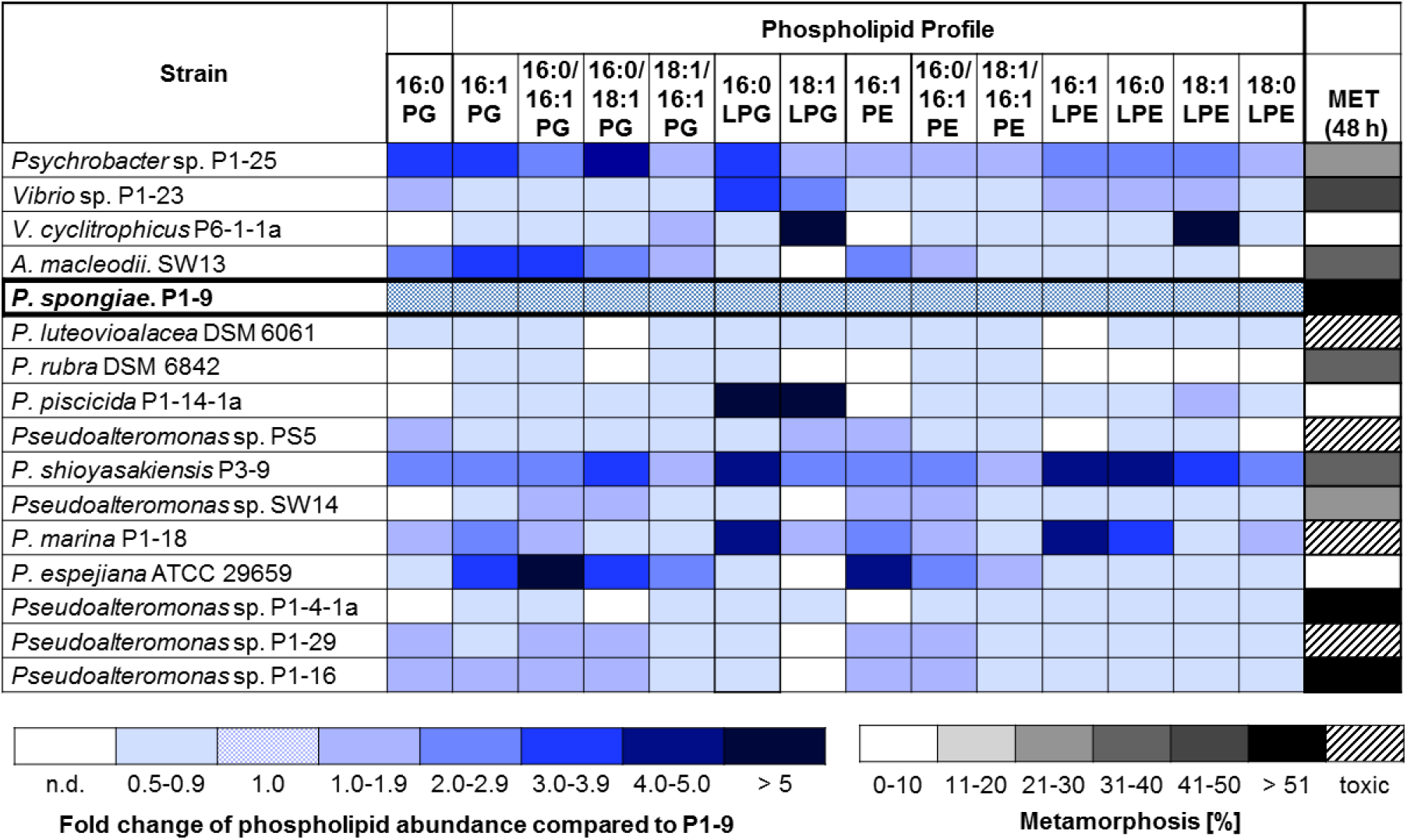
Relative phospholipid analysis of enriched cell extracts from different bacterial isolates (phospholipid abundance in *Pseudoalteromonas spongiae* P1-9 was set 1.0) and metamorphosis assay with bacterial lipid extracts (100 µg/mL). Transformation rates were evaluated after 48 h. Mean values of replicates are visualized as color-coded 10% step-gradient (n = 3, not shown: negative control: ASW; positive control: CsCl).

### Analysis of morphogenic polysaccharides

We then focused on the analysis of the most active HMW fractions (up to 80-100% metamorphosis) retrieved from aqueous extracts of bacterial biomass (Figure 2). First, the HMW fraction was purified using size-exclusion filtration (Figure S9) and fractions causing more than 20% of all larvae to metamorphose were analyzed by ^1^H NMR. Comparative analysis indicated towards a complex mixture of yet unknown polysaccharides (Figure S21) and further bioassay-guided purification using Sephadex G25 was performed. Again, NMR and HRMS analysis of the most active fraction resulted in the identification of a morphogenic polysaccharide consisting of repeating -(1’→4)-α-L-Rha-(1→3’)-D-Man-units (Table S2, Figure S19-S25). The rhamnose/mannose composition was confirmed by acid hydrolysis using 6 M HCl, followed by TMS derivatization and GC-MS analysis and comparison with commercial standards (Figure 16,). The purified polysaccharide, from now on named Rha-Man, showed a clear dose-dependent morphogenic induction of up to 80% transformation within 48 h (150 µg/mL, Figure S17). Partial acid hydrolysis resulted in the loss of morphogenic activity; similarly, rhamnose and glucose monomers showed no inducing effect.

Comparative NMR-analysis of bioactive HMW fractions clearly indicated that Rha-Man was not the only EPS within the bioactive fraction (Figure S19). However, due to the inherent difficulties associated with the separation and structural characterization of polysaccharides, we subsequently tested commercial and structurally defined bacterial polysaccharides (Figure 9). Intriguingly, curdlan, a well-known polysaccharide (50-200 kDa) with ß-1,3-glycosidic linkage and produced by the Gram-negative bacterium *Alcaligenes faecalis*, induced more than 50% of all larvae to undergo metamorphosis within 48 h. In contrast, paramylon, the 500 kDa derivative of curdlan, which differs in its average length and three-dimensional structure,^33^ and charged EPS derivatives, such as hyaluronic acid and heparin, did not induce metamorphosis. We then questioned, if combinations of EPS and LPS would result in the same morphogenic activities as observed for OMVs or biofilms, and indeed, combinations of curdlan (15 µg/mL each) with either 16:0 LPG, or 18:1 LPE (10 µg/mL each), or both, induced metamorphosis in more than 75% of all larvae causing the formation of a fully functional primary polyp within 24 h (Figure 9B).

**Figure 9.**
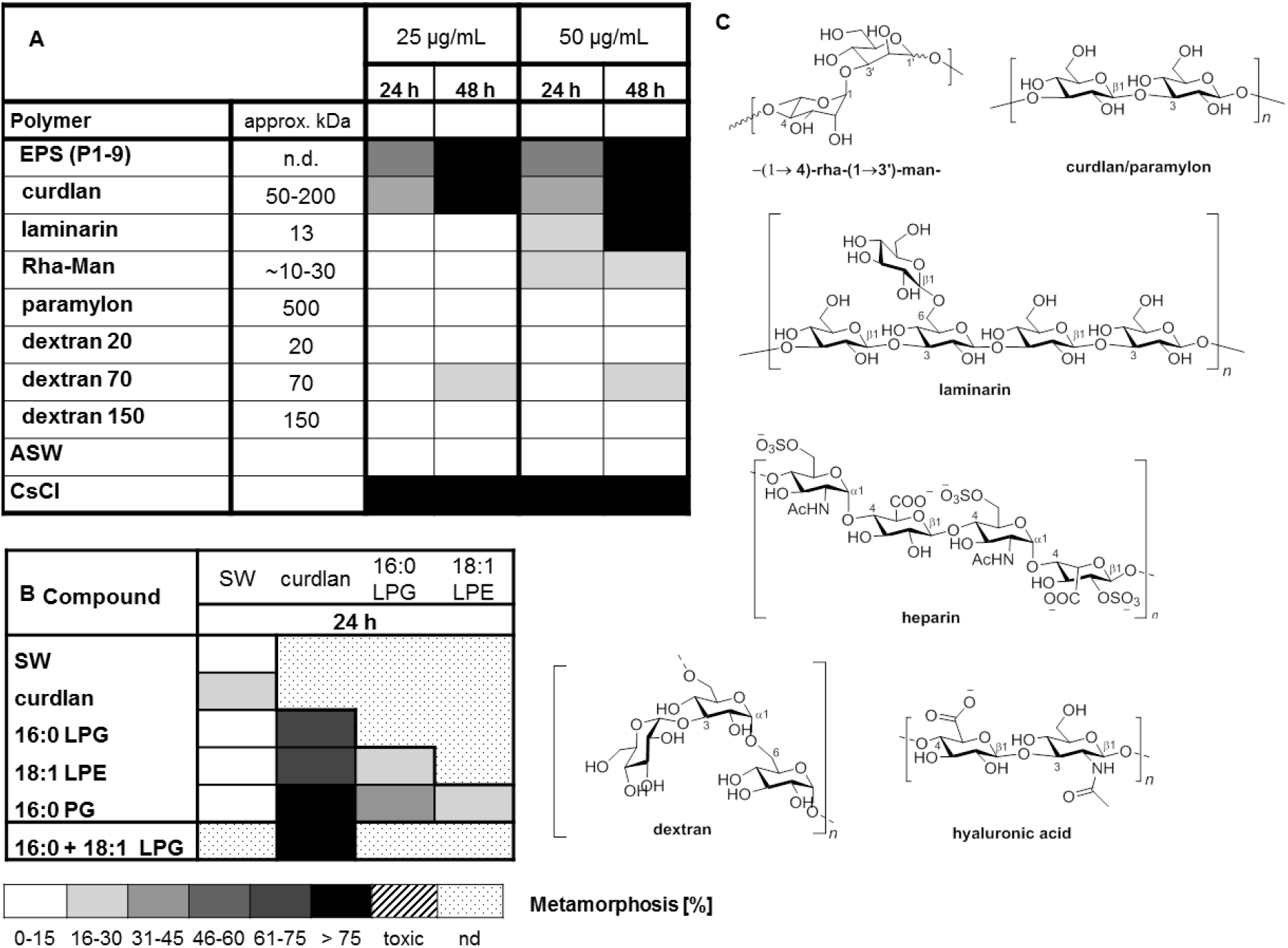
A) Metamorphosis assay with enriched and purified polysaccharides from P1-9 and commercial polysaccharides using (25 µg/mL) (n = 4); transformation rates were determined after 24 h and 48 h, respectively. B) Metamorphosis assay with commercial phospholipids (10 µg per lipid) and curdlan (15 µg) after 24 h. Mean values of replicates are visualized as color-coded 15% step-gradient (n = 3). C) Representative structures of repeating units of tested polysaccharides.

## Discussion

The broadly distributed marine hydroid polyp *H. echinata* serves as an established model system to study settlement and metamorphosis. While artificial induction with CsCl has long served as reliable and experimentally valuable method to induce metamorphosis in *H. echinata* in a synchronized way for downstream analyses, it does not enable anymore insights into the true induction of metamorphosis that is essential to the survival of the organisms. It was with this phenomenon that we asked the questions, which bacterial species and which bacterial metabolites are naturally abundant in marine biofilms and enable *H. echinata* to settle and metamorphose.

To determine the nature of the bioactive compound(s), we categorized a representative group of marine bacteria isolated from *Hydractina*, and commonly found in marine environments according to their ability to induce metamorphosis in a mono-species-based biofilm assay. Subsequent HRMS^2^-based analysis and bioassay-guided fractionation allowed the identification of two types of bacterial morphogens, a subset of (lyso)phospholipids and the exopolysaccharide Rha-Man from *Pseudoalteromonas* sp. P1-9 that induced the morphogenic transformation of *Hydractinia* planula larva into the primary polyp.

In case of phospholipids, in was notable that lysophospholipids 16:0/18:1 LPG, 18:0 LPE/PE and 16:0 LPA/PA induced metamorphosis as single compounds and in combination with other phospholipids, while other phospholipids were found to be inactive; however a clear structure-activity relation was not yet possible to deduce. Tracking of fluorescence-labled phospholipids revealed their cellular uptake, but a specific localization in different parts was not observed. These results in combination with the observation that phospholipid-enriched extracts from other bacterial strains induced metamorphosis, led to the assumption that the phospholipid composition of a bacterial biofilm dictates the observed morphogenic response and stimulating effects of single compounds might be overruled by the presence of toxic metabolites and/or inhibiting effects. In case of polysaccharides, a *Pseudoalteromonas* sp. P1-9 derived Rha-Man polysaccharide and *A. faecalis*-derived curdlan were found to induce the full transformation in a dose-response fashion with curdlan being the more active derivative. Intriguingly, combinations of phospholipids and curdlan induced the full morphogenic transition at rates exceeding in part the sum of single compound contributions and resulting in the complete transformation of almost all larvae within 24 h.

Although it is generally difficult to deduce whether the bioactive metabolites identified in the laboratory are the same as those that are responsible in Nature when larvae are exploring habitats, or if these signals only interact with downstream signaling cascades, the chemical nature and localization of the identified morphogens support their ecological relevance. As Gram-negative bacteria, including those from marine environment, produce OMVs,^13,25-27^ secrete specific EPS molecules to sustain the 3D integrity of the biofilm, ^34,35^ and release pieces of cells membrane when disrupted, both - phospholipids and EPS - would be the first bacterial molecules to interact with a larva upon physical contact with the biofilm matrix. For *Hydractinia* it has been proposed from early on that neurosensory cells dominantly located at or near the anterior pole and in part on its tapered posterior tip are responsible for the detection of the bacterial cues.^16,17^ Upon physical contact with lipid-rich vesicles and cell fragments naturally present within marine biofilms, phospholipids could passively integrate into the *Hydractinia* membrane; a hypothesis that is supported by our findings that fluorescence-labeled phospholipids are quickly integrated into membranes of *Hydractinia* larvae. Once integrated, specific phospholipids could act as ligands for intracellular receptors or induce changes in membrane fluidity resulting in the recruitment of, e.g., PKC involved in cellular signaling processes.^36,37,38,39^ In addition, LPAs have been recognized as potent mitogen in humans since decades due to their interactions with G-protein-coupled receptors (GPCRs), thereby altering many different cellular responses in humans, such as proliferation, survival, cytoskeletal changes or calcium influx.^40^ Thus, a homologous mode of action could be hypothesized for *Hydractinia*.

Similarly, polysaccharidesrequire the detection via dedicated receptors as described for EPS-mediated host-pathogen interactions^41,42^ and metamorphosis in other marine animals.^43^ In particular curdlan has been previously reported to act on lectin-type receptors in humans; a receptor-type which have also been detected in a *Hydractinia* transcriptome study.^44^ The structural variability of both compound classes and their variable abundance within the bacterial world, ^45,46,47,48^ raises the question on how *Hydractinia* selects for a suitable habitat to increase the likelihood of survival? Our study provides first evidence that the variation in bacterial induction results from additive or even synergistic effects of the identified bacterial signals and suggests that high percentages of larvae transformation are mostly achieved by a composition of these factors, and not only by the presence of single active molecules. Only few studies have so far focused on synergistic effects of morphogens in the marine environment, which includes studies on sulfonolipids (RIFs) and LPEs, that in combination induce the formation of predatory rosette-like stage of the choanoflagellate *S. rosetta*,^49,50,51^ and the recent identification of synergistically acting nucleobases from marine bacteria, that induce metamorphosis of the invasive fouling mussel *Mytilopsis sallei*.^52^

## Conclusion

The marine hydroid polyp *H. echinata* serves as an excellent model organism to study larvae settlement and metamorphosis in response to marine biofilm-dwelling bacteria.

We extended the studies on bacterial isolates that cause larvae of *H. echinata* to settle and metamorphose after contacting the bacterial biofilm and found that larvae settle in response to outer membrane vesicles and cell fragments of one of those inducing species. The identified bacterial metabolites satisfy the ecological criteria for acceptance as an inductive cue due to their abundance in bacterial biofilms, but raise now the question on how the selective settlement of *H. echinata* is achieved and the underlying molecule-receptor interactions. Results of our studies will allow us to manifest *Hydractinia* as a model system to investigate more closely host-microbe interactions and sheet light into the long-standing mystery on how bacterial signals trigger animal development in the marine world.

## Materials and Methods

Supplementary Information (SI) is available at https://zenodo.org with the DOI:10.5281/zenodo.4537693. SI material contains details on fermentation, cultivation, sample treatment, bioassays, isolation procedures, ESI-HRMS, ^1^H NMR, ^13^C NMR, and 2D NMR spectra as well as additional assay data.

### Cultivation of *Pseudoalteromonas* sp. P1-9

A preculture of marine bacteria were grown for 3 days at 30 °C (160 rpm) in MB medium and used as inoculum. For isolation of inducing signals, a 7 mL preculture of *Pseudoalteromonas* sp. P1-9 was used to inoculate 500 mL MB broth and incubated at 30 °C for three days (150 rpm). Then, cultures were centrifuged for 20 min at 4 °C (4000 rpm), and supernatant separated from cell pellet. Cell pellet was washed three times (suspended in 5 mL sterile PBS, centrifuged (20 min, 4 °C, 4000 rpm) and the supernatant was discarded), and then suspended in 1 mL sterile water and sonicated on ice (5 x for 30 sec with 30 sec break). To remove soluble cytosolic components, samples were centrifuged (10000 rpm, 10 min, 4 °C), the supernatant discarded and the resulting cell pellet lyophilized overnight (P1-9A samples). Lyophilized cell pellet was used for further enzymatic (DNase, RNase) and chemical treatments (see supporting information for details).

### *H. echinata* husbandary

*H. echinata* colonies were obtained as single colonies on gastropod shells from the Alfred Wegener Institute, Helmholtz Centre for Polar and Marine Research (Helgoland, Germany). Adult polyps on shells were kept in artificial seawater (salinity of 33.2– 33.7‰, pH 8.2–8.3 and 16 °C) in aerated tanks maintained in 16 h light/8 h dark cycle and were fed daily using 3-7 day old nauplii of *Artemia salina*. Fertilized eggs were collected in the two- to four-cell stage 3 hours after spawning event and transferred into freshly sterile seawater.

### Bioassay

After spawning event, fertilized eggs were collected and kept in sterile-filtered artificial sea water (ASW) at 18 °C. After two days, developing larvae were washed two-three times with sterile ASW and transferred to a new sterile petri dish filled with fresh ASW. The procedure was repeated every second day until use and larvae were kept at 20 °C. For bioassays, 20-30 competent larvae (kept in ASW) were added to a test well (sterile 24 well-plates) and well was filled up with ASW to reach 2 mL. For all experiments: negative control (larvae kept in sterile ASW) and positive control (larvae induced with a CsCl solution (6 mM final concentration)) were included. Experiments were performed at 20 °C as triplicates and repeated with larvae from at least two different spawning events.

### Colony-based assay

Bacterial strains were grown on a marine broth (MB) agar (Carl Roth) or LB agar for three days. A single colony was suspended in 100 µL PBS (OD600 of 0.2), and 5 µL of the bacterial suspension transferred to a sterile well (24-well plates). Cells were incubated for 20 min under laminar flow to allow for surface attachment, and then used for testing.

### Testing of biosamples and chemical extracts

Samples were centrifuged at 10000 rpm for 10 min at 4 °C resulting in an insoluble pellet (“pellet”) and supernatant (“sup”). For assays, the pellet was suspended in 1 mL sterile sea water and supernatant was used without further dilution. For testing, 60 µL of each sample were added to a sterile 24-well, dried under sterile laminar for 20 min, and then used for testing.

### Testing of phospholipids

Each (lyso)phospholipid was dissolved in MeOH to give a 1 mg/mL stock solution. The respective test amount was added to each well of a sterile well-plate (e.g. sterile 24-well plates). Samples were dried under sterile laminar flow and then used for testing.

### Testing of commercial polysaccharides

Exopolysaccharides were dissolved in ddH_2_O to give a 1.0 mg/mL stock solution and sterilized by membrane filtration (0.22 µm). Insoluble polysaccharide, eg. curdlan and paramylon, were autoclaved at 120 °C for 10 min. For assays, 100 µL or 50 µL of each stock solution were applied per well, dried under sterile laminar flow and then used for testing.

## Structure elucidation

### Analysis of phospholipid content

Bacteria were cultured in 50 mL MB (180 rpm, 30 °C, 2 d) and supernatant was separated by centrifugation (4000 g, r.t. 20 min) and discharged The cell pellet was collected, lyophilized and extracted twice with MeOH (10 mL) assisted by ultrasonication. Methanolic extracts were dried, re-dissolved into MeOH to yield a 5 mg/mL stock solution for testing and LC-HRMS analysis.

### HPLC-HRMS analysis

Measurement were performed on a Thermo Q-Exactive Plus using the following gradient: 0–1 min, 20% B; 1–2 min, 20%–40% B; 2–25 min, 40%–92.5% B; 25–26 min, 92.5%–100% B; 26–35 min, 100% B; 35–35.1 min, 100%–20% B; 35.1–38 min, 20% B (A: dd H_2_O with 20 mM ammonium formate pH 4.5; B: 50% *i*PrOH/50% MeCN), flow rate: 0.3 mL/min. Metabolite separation was followed by a data-dependent MS/MS analysis in both positive (MS^1^) and negative (MS^1^ and MS^2^) ionization modes.

### GNPS mediated molecules networking

Metabolomic raw data files were recorded on Thermo QExactive MS instrument and firstly converted to .mzXML by ProteoWizard, then uploaded to GNPS (Global Natural Products Social Molecular Networking) server.

### Purification of phospholipids

Lyophilized cell pellet derived from 1 L culture (MB, 30 °C, two days, 120 rpm) was extracted with 500 mL MeOH. Insoluble cell debris was filtered off and the resulting methanolic filtrate collected and concentrated under reduced pressure. The extract was loaded on a SiOH SPE cartridge (2 g) and metabolites eluted using a step gradient of solvent mixtures (cyclohexane, EtOAC and MeOH). Metabolites were collected in eight fractions and the solvent was removed under reduced pressure. The metabolite content of each fraction was analyzed by UHPLC-MS and NMR and the morphogenic properties evaluated.

### Purification of polysaccharide

Lyophilized cell pellet was extracted with ddH2O and the resulting crude extract (170 mg) purified by size exclusion chromatography by Sephadex G25. Active fraction (orange band (53 mg)) was again lyophilized, suspended into 1 mL H_2_O, heated to 95 °C for 1 h, then cooled on ice for 5 min and submitted to DNAse, RNAse and proteinase K treatments. The resulting solution was centrifuged (13,000 rpm, 20 min), filtered through 0.45 µm subjected to semipreparative size exclusion chromatography (Shodex GF310HQ column). Collected fractions were lyophilized and submitted to the metamorphosis assay and NMR analysis.

### Scanning electron microscopy (SEM)

Bacteria were fixed with glutaraldehyde (2.5%, v/v) for 1h at RT while sedimenting on poly-L-lysine coated coverslips. After two washings in cacodylate buffer (100 mM, Ph 7.2) cells were dehydrated by increasing concentrations of ethanol, followed by critical point drying in a Leica EM CPD300 Automated Critical Point Dryer (Leica, Germany). Then, the cover slips were mounted on aluminum sample holders (stubs) and gold sputter coated (layer thickness 20 nm) in a BAL-TEC SCD 005 Sputter Coater (BAL-TEC, Liechtenstein). Images were acquired with a Zeiss (LEO) 1530 Gemini field emission scanning electron microscope (Zeiss AG, Germany) at 4 kV acceleration voltage and a working distance of 3-4 mm using an Inlense secondary electron detector.

### Cryo-Transmission Electron Microscopy (TEM)

A drop of a freshly prepared OMV solution (2 μl) was placed on a R3.5/1 holey carbon filmed copper grid (Quantifoil Micro Tools GmbH, Germany). The sample was rapidly plunge-frozen in liquid ethane at - 180°C. The frozen grid was transferred into a liquid nitrogen cooled Gatan 626-DH cryo-holder (Gatan Inc., USA) and inserted into a Philips CM 120 cryo-TEM (Philips, Netherlands) operated at 120 kV accelerating voltage. Images were acquired with a 1k × 1k FastScan-F114 CCD-camera (TVIPS GmbH, Germany). Isolated OMV/minicell solution (20 µl) were placed for 1 min onto hydrophilic, Formvar®/carbon-filmed copper grids (Quantifoil Micro Tools GmbH, Germany), washed twice on drops of distilled water, and stained on a drop of 2 % uranyl acetate in distilled water. Samples were imaged in a Zeiss EM902A electron microscope (Carl Zeiss AG, Germany) operated at 80 kV accelerating voltage. Images were acquired with a 1k × 1k FastScan-F114 CCD-camera (TVIPS GmbH, Germany).

## Acknowledgments

We are grateful for financial support from the German Research Foundation (DFG, BE 4799/2-1). CB greatly acknowledges funding by the ERC (ERC Grant number: 802736, MORPHEUS). MR is generously supported by the Jena School for Microbial Communication (JSMC, DFG). Mrs. Heike Heinecke (HKI) for recording NMR spectra, Mrs. Andrea Perner (HKI) for HRMS measurements, Toni Neuwirth for GC-MS measurements and Johan Kufs for assisting in fluorescence imaging. We also would like to thank Theresa Jautzus for critical comments on the manuscript. We are very grateful for the support by the Alfred-Wegner Sylt team and Ms Birgit Hussel for their hospitality during field work.

